# Stress response of fire salamander larvae differs between habitat types

**DOI:** 10.1101/2023.08.07.552279

**Authors:** Laura Schulte, Pia Oswald, Max Mühlenhaupt, Edith Ossendorf, Sabine Kruse, Sylvia Kaiser, Barbara A. Caspers

## Abstract

Different habitats can impose varying selection pressures on individuals of the same species. Larvae of the European fire salamander (*Salamandra salamandra*) can inhabit two different habitats: streams and ponds. Streams are characterised by lower predation risk and intraspecific density and higher food availability than ponds. Thus, ponds are considered a less suitable habitat. To investigate the differential impacts of the two habitats on larval physiology, we measured the stress response of larvae living in the two different habitats. After successfully validating the measure of water-borne corticosterone (CORT) concentrations in fire salamander larvae, we measured the baseline and stress-induced CORT of 64 larvae from two pond and two stream locations in the field. We found larvae in ponds to be more stressed than larvae in streams. Additionally, we performed a reciprocal transplant experiment and tested whether larvae can adapt their stress response to changing habitat conditions. After two weeks of transfer, we did not find an increase in CORT when comparing stress-induced CORT values with baseline CORT values in larvae transferred into ponds. However, larvae transferred into streams exhibited an increase in the stress-induced CORT response. Moreover, transfer into ponds as well as larvae originating from ponds showed reduced growth rates, indicating that ponds might be the more stressful habitat, as it negatively affected larval physiology. These results show that noninvasive hormone measurements can provide information on habitat quality and potential adaptation and thus emphasis the potential for its use in conservation efforts.

## Introduction

Individuals of the same species often reside in different habitats, which can come with different selection pressures. For instance, the common raven (*Corvus corax*) inhabits different habitat types and adapts its diet to the food resources available in each type of habitat (Harju et al., 2021). Other species show adaptive morphological changes in response to different habitats, such as the Mexican tetra (*Astyanax mexicanus*) (Jeffery, 2020). This fish species is found in both surface and cave waters and has adapted to very different environments, such as reduced or no eyes in the cave type. Selection pressures might also shift due to environmental changes, such as human-induced climate change. For example, precipitation becomes unpredictable and unstable, causing droughts and flooding. These extreme weather events will likely occur more frequently in the future and can substantially impact aquatic ecosystems in particular (Bickford et al., 2018; Blaustein et al., 2001; IPCC, 2021). Stagnant water bodies, for example, are expected to dry out and become less abundant in the future due to droughts, while floods might lead to overflooding of streams, causing mass-drift events (Bickford et al., 2018). Therefore, conforming to a changing environment imposes great challenges organisms have to face, both now and in the future.

Responding to a changing environment is particularly challenging for amphibians because most of them rely on water bodies for reproduction and they already show strong population declines due to human-induced environmental changes (Bickford et al., 2018; Blaustein et al., 2001; Huey et al., 2012; Woodley, 2018). Environmental changes can influence habitat quality in different ways. However, knowledge regarding which habitat type is less stressful for a species is often missing. Environmental stressors have the potential to increase the level of glucocorticoids (Denver, 2009a; Rollins-Smith, 2001; Sapolsky et al., 2000). Hence, one way to evaluate stress responses in vertebrates is to measure the release of glucocorticoids. Glucocorticoid (GC) hormones, such as cortisol and corticosterone (CORT), are useful biomarkers to assess the condition of amphibians. These “stress hormones” are responsible for homeostasis, growth levels, and control immune functions, and CORT is the most responsive stress hormone in amphibians (Idler, 1972). Stress hormones are responsible for changes in morphology, physiology and behaviour and can also disrupt the amphibian life cycle (Bókony et al., 2021; Denver, 2009b; Idler, 1972; McEwen and Wingfield, 2003; Middlemis Maher et al., 2013). Long-term stress can be assessed by measuring the baseline CORT release, while the stress response towards an acute stressor can be assessed by measuring stress-induced CORT release (Narayan et al., 2019; Vitousek et al., 2019). When stress hormones are chronically elevated, they can negatively impact organisms, such as higher disease rates (Cyr and Romero, 2009; Forsburg et al., 2019; McEwen and Wingfield, 2003; Paul et al., 2022). However, when they are acute, they can have positive short-term effects, such as increased immune response and survival (Sapolsky et al., 2000).

In general, amphibian larvae show great phenotypic plasticity towards abiotic and biotic factors in their habitat (Denver, 1997). Organisms that show high levels of plasticity will possibly be able to respond faster to a changing and unstable habitat (Newman, 1992). Habitat conditions can influence the endocrine system and the endocrine system in turn has a major impact on the plasticity of individuals. Thus, the larval habitat, and or different larval habitats, respectively, can indirectly influence on fitness via the endocrine system (De Witt and Scheiner, 2005). Hence, investigating the individual baseline and stress induced CORT release will provide insights about potential long-term environmental stressors and the ability of individuals to cope with acute stress.

Larvae of the European fire salamander (*Salamandra salamandra*) occur in very different habitats. Larvae are deposited mainly into first-order streams (Thiesmeier, 2004) but also into ephemeral ponds (Steinfartz et al., 2007; Weitere et al., 2004). These habitats differ in several aspects. Salamander larvae in streams face little predation pressure, for example, from dragonfly larvae (Giesenberg, 1991), while the predation pressure in ponds is generally higher, because of newt species for instance (Thiesmeier, 2004; Weitere et al., 2004). Analyses of potential food resources for fire salamander larvae have shown that streams offer greater energetic value of potential food than ponds (Reinhardt et al., 2013). Additionally, the densities of conspecifics in ponds are much higher than those in streams (Weitere et al., 2004). It can therefore be assumed that ponds are a more stressful habitat for fire salamander larvae. Laboratory studies found that the amount of food available to larvae influences risk-taking behaviour (Krause et al., 2011) and the amount of yellow colouration after metamorphosis (Caspers et al., 2020), both having potential fitness consequences. Fire salamander larvae also differ in their risk-taking behaviour depending on their habitat type (Oswald et al., 2020). The larvae, however, not only differ in behaviour but also in their morphology, as larvae from ponds have a higher tail fin depth (Schulte, 2008). Additionally, a previous reciprocal transplant experiment (RTE) showed that larvae in pond habitats perform better than larvae in stream habitats regarding their growth rate and survival after metamorphosis (Oswald and Caspers, 2022). In summary, it is not yet understood which larval habitat reflects less stressful conditions for the larvae: ponds or streams.

The larval stage represents the most critical time during the fire salamander life cycle, as fire salamanders rely on aquatic habitats for reproduction (except for some Iberian subspecies, see Thiesmeier, 2004). Metamorphosed salamanders have almost no natural predators and live in more stable habitats (Thiesmeier, 2004). Thus, the larval stage should be the focus of conservation measures. Considering the environmental changes caused by global warming and extreme weather events, it is important to understand which larval habitat is more stressful for fire salamanders in light of conservation physiology.

With this study, we aimed to quantify how larvae from two habitat types (ponds and streams) differ in their stress physiology. Additionally, we performed a reciprocal transplant experiment (RTE) in which we transferred larvae between these two habitat types. Thus, we assessed whether larvae could adapt to changing habitat conditions. We predicted to find i) differences in the stress responses between pond and stream larvae, and we expected larvae in ponds to have higher baseline CORT release levels than stream larvae due to the harsher conditions in ponds. Furthermore, we wanted to test whether larvae are plastic in their stress response and can conform to the given conditions of a specific habitat. If this is the case, we predicted to find (ii) that the transfer habitat influences the stress response, whereas if larvae cannot conform to the given habitat, we expected to see an effect of the original habitat after the RTE.

## Methods

### Study area

The study sites are located in the Kottenforst, Bonn, Germany (50°39’30.58” N, 7°4’4.70” E, figure 1). The forest mostly consists of *Luzulo-Fagetum* and *Stellario-Carpinetum* communities with European hornbeam (*Carpinus betulus*), common oak (*Quercus robur*) and European beech (*Fagus sylvatica*) as the main tree species forming an ideal habitat for fire salamanders (Thiesmeier, 2004). In some parts of the forest, however, Scotch pine (*Pinus sylvestris*) and European spruce (*Picea abies*) dominate the landscape, while many European spruce trees have died in recent years due to beetle infestations. The Kottenforst holds several streams but also contemporary ponds that are both used for larval deposition by fire salamander females (Hendrix et al., 2017; Reinhardt, 2014; Steinfartz et al., 2007; Weitere et al., 2004). These two different habitat types are associated with two different genotypes of the fire salamander, indicating a recent local adaptation (Steinfartz et al., 2007). A common garden experiment revealed different larval deposition strategies in female fire salamanders of the two habitat types (Caspers et al., 2015).

**Figure 1:**
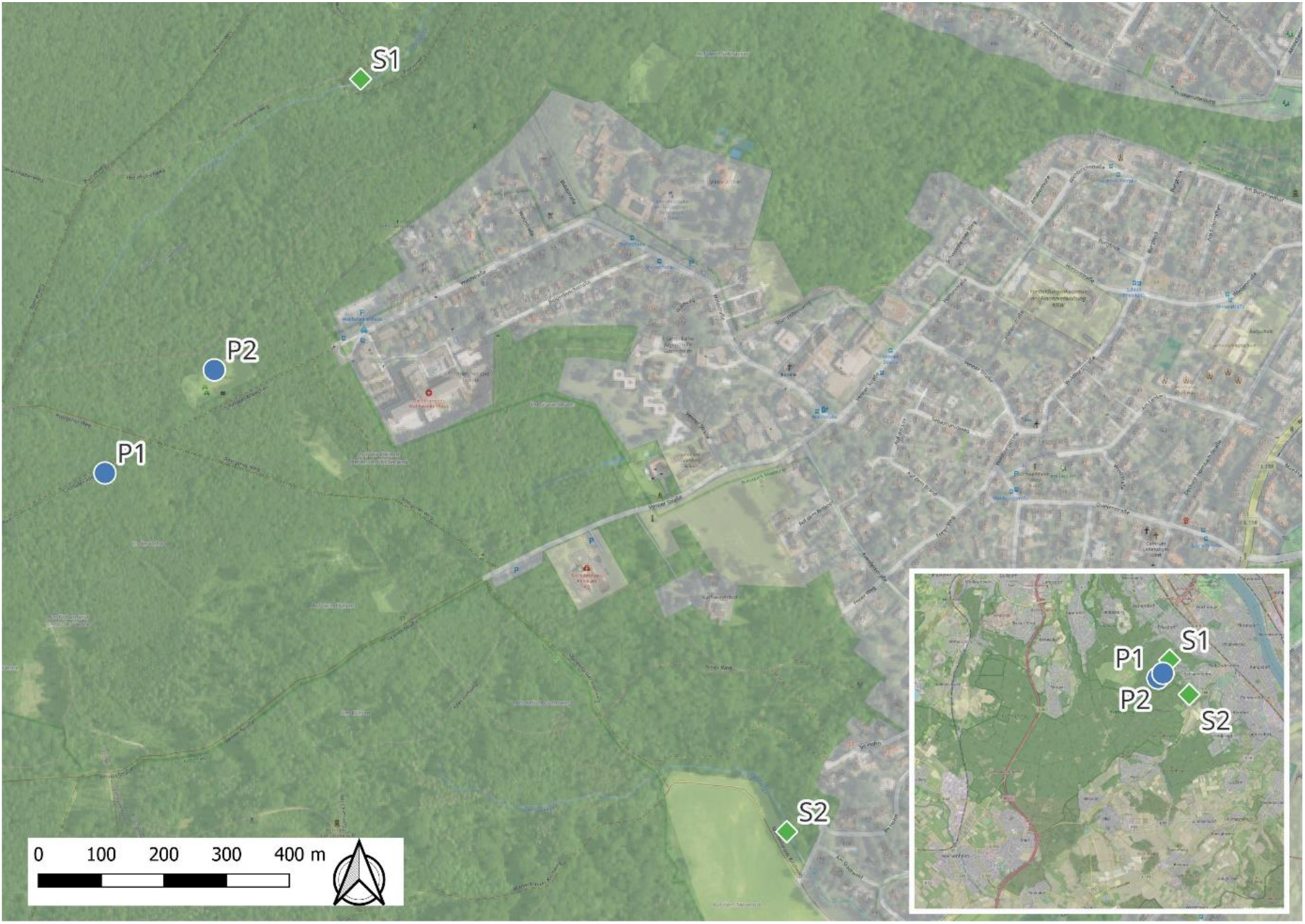
The four study locations in the Kottenforst, Bonn, Germany. The two blue dots represent the two pond locations (P1, P2), and the green squares represent the two stream locations (S1, S2). The inlet provides a larger overview of the Kottenforst.

### Water-borne hormone sampling

In April 2022, we tested 64 fire salamander larvae for baseline and stress-induced CORT release using water-borne hormone sampling. Waterborne-hormone sampling is a noninvasive and nonlethal method to assess an organism’s stress response (Forsburg et al., 2019; Gabor et al., 2013). In the past, CORT levels were often measured by euthanising the animal, especially small organisms (Gabor et al., 2013; Millikin et al., 2019). Thus, measuring water-borne hormone release is a promising alternative and particularly beneficial for small aquatic or semiterrestrial organisms (Narayan et al., 2019). Moreover, it allows repeated sampling (Gabor et al., 2013; Narayan et al., 2019), providing the opportunity to study individual conditions over a longer period or under changing conditions. We captured larvae from 4 locations, two ponds and two streams (N = 16 from each location, figure 1). None of the captured larvae showed signs of metamorphosis (e.g., reduced gills) or were morphologically impaired (e.g., missing tail, missing limbs); consequently, all the captured larvae were used in the study. Immediately after capture, we placed the larvae individually into a plastic jar with 40 mL of tap water each (figure 2). The tap water was identical for all sampling locations. Each larva was kept in the jar for 60 min to measure the baseline CORT release rate per hour (altered after Bryant et al., 2022; Charbonnier et al., 2018; Gabor et al., 2015, 2013, figure 2).

**Figure 2:**
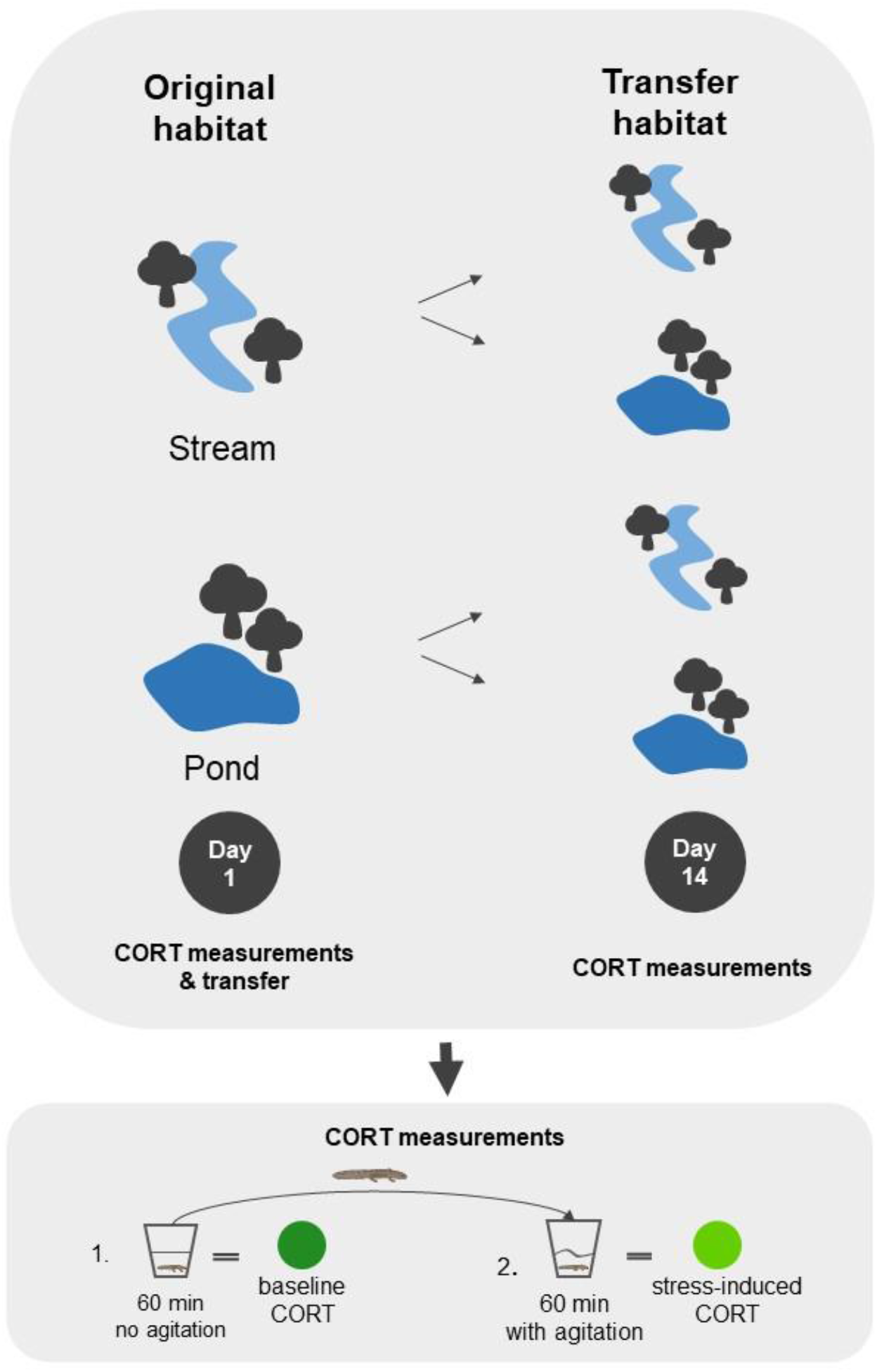
Top: On day 1 of the experiment, larvae were tested for baseline (60 min without agitation) and stress-induced (60 minutes with agitation) corticosterone (CORT) immediately after capturing them from their original habitat. Afterwards, they were transferred reciprocally between four locations (two ponds and two streams, including the location of origin). On day 14 after the transfer, larvae were measured again for baseline and stress-induced CORT, this time at the transfer location. Bottom: For the CORT measurements, larvae were first tested for baseline CORT. We took the larva from the water body with a dip net. We immediately placed them in a plastic jar with 40 mL of tap water and kept them there for 60 minutes. Afterwards, we placed them in 40 mL of fresh tap water to assess the stress-induced CORT level. Here, we again kept them for 60 minutes, but this time we agitated the jars for one minute every three minutes.

The jars were placed in the shadow and left undisturbed for the first 60 minutes. After 60 minutes, we transferred each larva into a new plastic jar with 40 mL of fresh tap water. There, we kept them for another 60 min and agitated them manually for one minute every three minutes to measure stress-induced CORT release (Bryant et al., 2022). The water from the first and second plastic jars was transferred separately into 50 ml Falcon tubes and stored in a portable car freezer (-10°C) immediately after conducting the experiment in the field (approximately 2.5 hours after starting with the baseline CORT). After returning from the field, we stored the samples at -20°C until further processing. We cleaned all plastic jars with ethanol and rinsed them with the same tap water used for the experiment (Charbonnier et al., 2018). Sampling was conducted during different times of the day, and we alternated the time of the experiments between the two habitats to control for potential circadian variation in CORT release levels. We also collected negative and positive hormone control samples to assess the quality of the assay (see below). Negative control samples consisted of 40 ml of tap water, which was the same tap water that we used for CORT measurements. They were also frozen at -20°C until the waterborne corticosterone concentrations were determined. We produced two different positive control samples by pooling samples of each of the 21 larvae. The positive control samples were aliquoted into 40 ml vials and kept at −20°C. We wore nitrile gloves during the entire experiment. After the CORT release rate measurement, we photographed the larvae from the right body side and from the top, and we measured the total length in mm (size). We sterilised all our equipment with *Virkon S* (LANXESS Biosecurity) before and after sampling to avoid the potential spread of the chytrid fungus *Batrachochytrium salamandrivorans* in our study area.

### Reciprocal transplant

To investigate whether the larvae can conform their CORT release rate to the changed habitat, we performed a reciprocal transplant experiment (following Sabino-Pinto and colleagues (2019) and Bletz and colleagues (2019)). Briefly, within our two habitat types (pond and stream) we used four locations (two streams and two ponds), resulting in the following treatment groups: stream-origin transferred into streams (St/St), stream-origin transferred into ponds (St/P), pond-origin transferred into ponds (P/P) and pond-origin transferred into streams (P/St) (figure 2).

To control for location and transfer effects, we transferred four larvae from each location into the same habitat type but into the different location, four from each location to each of the two locations of the other habitat type, and four remained in their original habitat (same location). After performing the measurements as described above (waterborne hormone sampling), we placed the larvae into individual enclosures (8.5 x 8.5 x 17 cm, similar to Bletz et al. 2016) and assigned them randomly to one of the treatment groups and locations. We kept the larvae for two weeks in the treatment and then repeated the CORT release measurement as described above (figure 2). The individual enclosures had small grits that allowed food items to enter the enclosure and prevented the larva from escaping. Additionally, they had styrofoam pieces attached to the top to enable floating and we equipped the enclosures with a stick to prevent drowning in case the larvae underwent metamorphosis. We checked the larvae every day to see if they were alive or about to undergo metamorphosis. All larvae were released after the two weeks of the experiment.

### Water-borne hormone extraction and determination of corticosterone

After thawing, the water-borne hormone samples were centrifuged (5 min 2500 x g). Hormones were then extracted from the supernatant under vacuum pressure using a Chromabond SPE vacuum chamber and HR-X SPE (solid phase extraction) columns (CHROMABOND^R^ HRX polyprobylene columns, 3 ml/60 mg/45 µm; Macherey-Nagel [# 730936P45], Düren, Germany). The columns were activated with 3 ml methanol (p.a.) and 3 ml demineralised water. After activation, columns were filled with 2 ml demineralised water. The samples were then added to the column using a 70 ml reservoir tube connected to the columns. The vacuum was regulated in such a way that samples were adsorbed to the HR-X columns at a flow rate ≤ 3 ml/min. After running the samples, the columns were first washed under maximum vacuum pressure with a column volume (i.e., 3 ml) of demineralised water, then pulled as dry as possible, sealed with Parafilm, and stored temporarily at -20°C. After thawing the columns, elution of corticosterone was performed with 1 ml methanol (p.a.) and an elution rate ≤ 1 ml/min. The methanol was evaporated directly afterwards in a vacuum concentrator, and corticosterone was resuspended in 250 µl ELISA buffer (Cayman Chemicals Inc. [# 501320]), and the samples were stored at -20°C until ELISA was performed. Corticosterone concentrations were determined in duplicate and according to the manufacturer’s instructions. Each ELISA plate was randomly loaded with samples from 2 larvae from each area (stream-stream, stream-pond, pond-pond, pond-stream). In addition to these 32 samples, two positive and four negative controls (two samples of tap water original habitat, two samples of tap water transfer habitat) were added per plate.

### Assay quality control

#### Cross reactions and inter- and intra-assay variation

Two positive control samples were run in duplicate on a total of six and eight plates to assess inter-assay variation, which was on average 7.7%. Furthermore, four pools of different hormone concentrations were used to determine intra-assay variance: corticosterone concentrations of each pool were determined 12 times for one plate, resulting in an average intra-assay coefficient of 6.6%. Finally, all samples were determined in duplicate, and determination of the sample was repeated if CV was larger than 10%. The antibody showed the following cross-reactivities: corticosterone 100%, 11-deoxycorticosterone 15.8%, prednisolone 3.4%, 11-dehydrocorticosterone 2.9%, cortisol 2.5%, progesterone 1.4%, aldosterone 0.47%, 17α-hydroxyprogesterone 0.21%, 11-deoxycortisol 0.14%, androstendione 0.11% and all other tested steroids < 0.1%.

### Linearity/Parallelism

To assess linearity, we ran a serial dilution of three pooled samples in duplicate (see Supplements table 1).

### Recovery rate

Two different recovery rates were determined:

1. After extracting corticosterone from the water, samples were resuspended in ELISA buffer (see above). Three pooled samples of different concentrations resulting from these resuspended samples were used to calculate the recovery rate of the assay. We conducted spikes by mixing equal volumes of the respective pooled sample with three different standards, which were also used for the standard curve in the ELISA. In addition, we determined the concentrations of the non-spiked and non-diluted as well as 1:2 diluted pooled control samples. All samples were determined in duplicate. The expected recovery concentrations were based on the known amount of corticosterone concentrations in the control samples (see Supplements table 2).
2. To assess the recovery rate of the HR-X SPE columns, we used the following procedure: four different standards, which were also used for the standard curve in the ELISA, were added to tap water, and the same procedure as described above was conducted. Furthermore, we prepared water samples containing each of four different standards and a pooled sample. The concentrations of the pooled sample were determined in the same assay. All samples were determined in duplicate. The expected recovery concentrations were based on the known amount of the standards and pooled samples minus the concentration of the negative controls (see Supplements table 3).

### Calculating corticosterone concentrations

The concentrations of corticosterone in pg/ml measured in the ELISA were multiplied by 0.25 ml (the volume of ELISA buffer used to resuspend the sample) after concentrations of negative control samples were subtracted from the sample values (a standard procedure that must be used because of matrix effects; Milikin et al., 2019). These values were standardised by dividing by the body size of each individual. Thus, corticosterone concentrations were given as “pg/size/h”.

### Statistical Analyses

The CORT measurements were first standardised to the size as pg/size/h (Gabor et al., 2013). We tested for normality with the Shapiro-Wilk test and homogeneity of variance using the variance test (*dplyr* R package). If normality was not given, we used the *best Normalise* R package to find the best transformation procedure of our data to fulfil the normalisation assumptions. For linear models (LM) and linear mixed effect models (LMM), we used the R packages *lme4* and *lmerTest* (Bates et al., 2015; Kuznetsova et al., 2017).

The first sampling point, i.e., before the reciprocal transplant experiment, aimed to test for a potential impact of the habitat on baseline CORT release and stress-induced CORT release. Therefore, we performed a linear mixed effect model with CORT release as the dependent variable and CORT type (baseline CORT or stress-induced CORT) and habitat (pond and stream) and the interaction between the two as fixed factors. We controlled for repeated measurements and potential location effects (e.g., sibling effect) by having individual ID and sampling location as random factors.

Similarly, we performed another LMM to investigate if there was an impact of the habitat on the baseline and stress-induced CORT release rate, this time after the reciprocal transplant experiment. We included both the habitat of origin and the transfer habitat in the model as well as the CORT type as fixed factors and the interaction between CORT type and transfer habitat and CORT type and habitat of origin. We used the ID and transfer location as random factors to control for repeated measurements as well as different habitat conditions in the transfer location.

To test for habitat-specific differences between baseline and stress-induced CORT, we performed two LMMs, one for ponds and one for streams both after the transfer, to test for differences in CORT between the CORT type (baseline and stress-induced CORT) as fixed factors and ID and location as random factors. To investigate whether pond and stream larvae differ in their baseline or stress-induced CORT, we compared only baseline CORT between ponds and streams and only stress-induced CORT between ponds and streams after the transfer. In addition, we compared the deviation in baseline and stress-induced CORT (=stress-induced CORT - baseline CORT) after the transfer per treatment group (P/P, P/St, St/P, St/St), and we performed a LM to test against the treatment groups. As a post hoc test, we used the package *emmeans* to compare the different treatment groups. Finally, we used another LM to investigate differences in the absolute growth rate between the treatment groups (P/P, P/S, St/P, St/St) after the transfer using *emmeans*. To test if there was a difference in the size of the larvae from the two habitat types before the transfer, we performed a t test. We conducted all statistical analyses with R Studio (version R 2022.02.3). We set the significance level to α = 0.05. No data were excluded, but three CORT measurements (1x baseline and 1 x stress-induced CORT release rate from one pond larva before transfer, 1 x stress-induced CORT release rate from one stream larva after being transferred into a pond) were lost due to handling error. All graphs were created using the package *ggpubr* (Kassambara, 2023).

### Limitations of the study

Newcomb Homan and colleagues (2003) found a different stress response in adult female and male spotted salamanders (*Ambystoma maculatum*). For fire salamander larvae, it is not possible to determine sex, and sex ratios in wild populations are unknown, which is why we cannot control for a potential sex effect. Other studies, however, have not found a sex effect on the stress response in a wild population of adult San Marcos salamander (*Eurycea nana)* (Gabor et al., 2016).

## Results

In total, we measured the CORT release rate of 64 larvae originating from four different locations (two ponds, two streams) before and after a reciprocal transplant experiment to test for a potential impact of the habitat (pond or stream) on larval CORT release rates. At both timepoints, we measured the baseline CORT release rates and stress-induced CORT release rates i) to test for a potential impact of the larval habitat and ii) to test whether larvae can conform to habitat-specific differences after being transferred.

### Before the transfer

We found no difference in the size of the larvae between the pond and stream habitats of origin (t test, p = 0.738). The stress-induced CORT release rate was significantly higher than the baseline CORT (p < 0.001, table 4) independent of the habitat (figure 3). Additionally, overall CORT release rates (baseline and stress-induced CORT) were significantly lower in streams than in ponds (p = 0.01). We did not find an interaction effect between the habitat type (pond or stream) and the CORT type (baseline or stress-induced CORT, p = 0.096) (figure 3).

**Figure 3:**
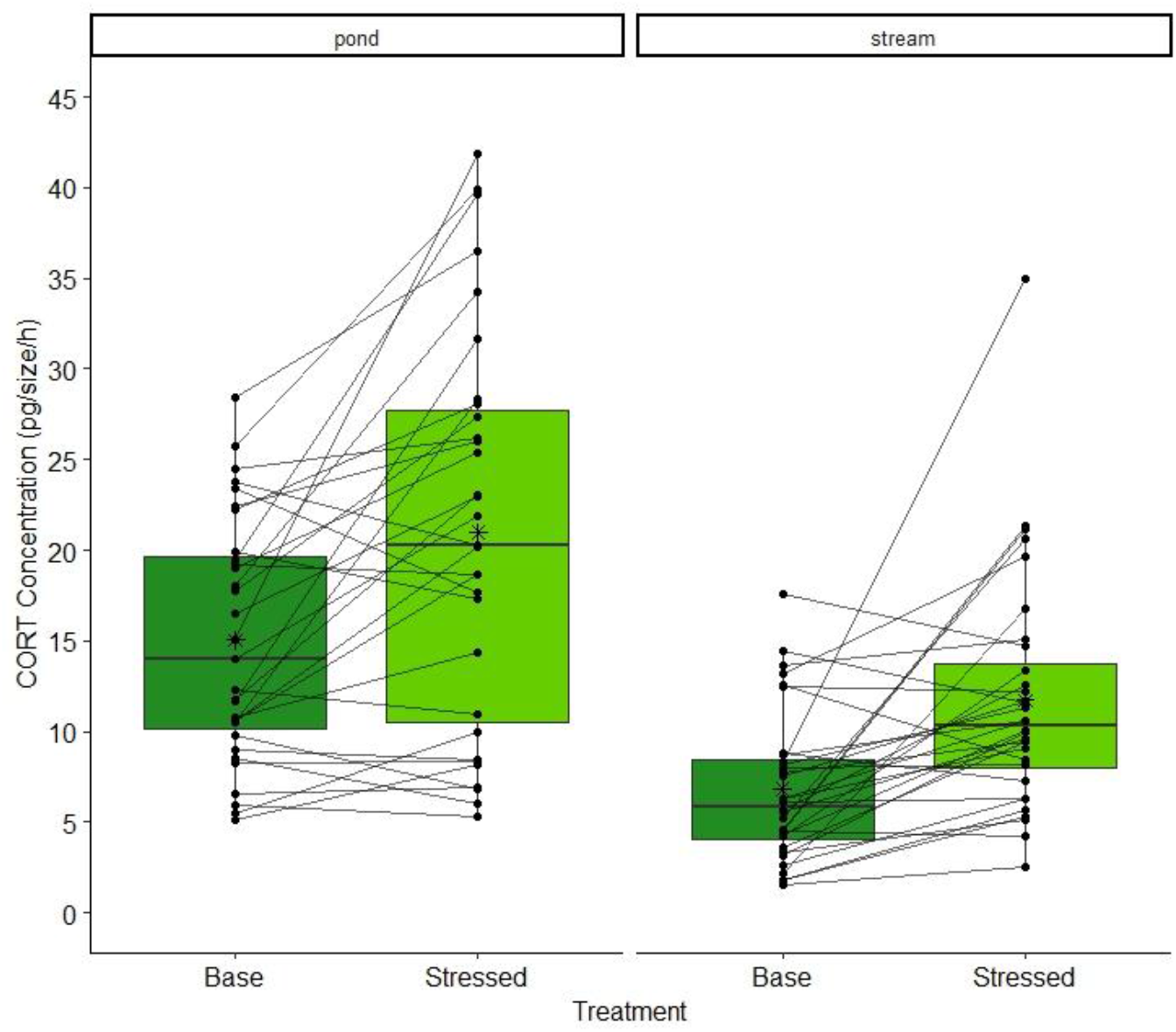
Waterborne corticosterone (CORT) concentration (pg/size/h) in fire salamander larvae (*Salamandra salamandra*). Shown are the baseline (Base) and stress-induced (Stressed) CORT release rates of pond (baseline N=31, stress-induced N=31) and stream (baseline N=32, stress-induced N=32) larvae before the transfer. Each point represents a single CORT measurement, and each line connects the two measurements per individual. The horizontal line represents the median, and the asterisks in the box represent the mean.

**Table 4:** Linear mixed effect model (LMM) for the corticosterone (CORT) release rate regarding the CORT type (baseline or stress-induced) before the transfer, habitat of origin, water temperature, interaction between CORT type and habitat of origin, ID and location. Significant effects are presented in bold.

| CORT release rate<br>(pg/size/h) | Dependent variable |  | Fixed effect |  |
| --- | --- | --- | --- | --- |
|  | Intercept | Estimate | Std Error | p value |
|  | **Stress-induced CORT** | **0.467** | **0.134** | **0.001 \*\*\*** |
|  | **Habitat stream** | **-1.159** | **0.211** | **0.01 \*** |
|  | Interaction Stress-induced<br>CORT & Habitat stream | 0.312 | 0.189 | 0.096 |

### After the transfer

We found a significant interaction between the treatment (i.e., baseline or stress-induced CORT) and the original habitat (table 5, p = 0.041). While larvae originating from streams did not show an increase in stress-induced CORT after the transfer period, larvae originating from ponds did (figure 4). Furthermore, we found an interaction between the treatment (i.e., baseline or stress-induced CORT) and the transfer habitat (table 5, p = 0.005), with larvae transferred into streams showing a significant increase in stress-induced CORT, while larvae transferred into ponds did not show this increase (figure 4).

**Figure 4:**
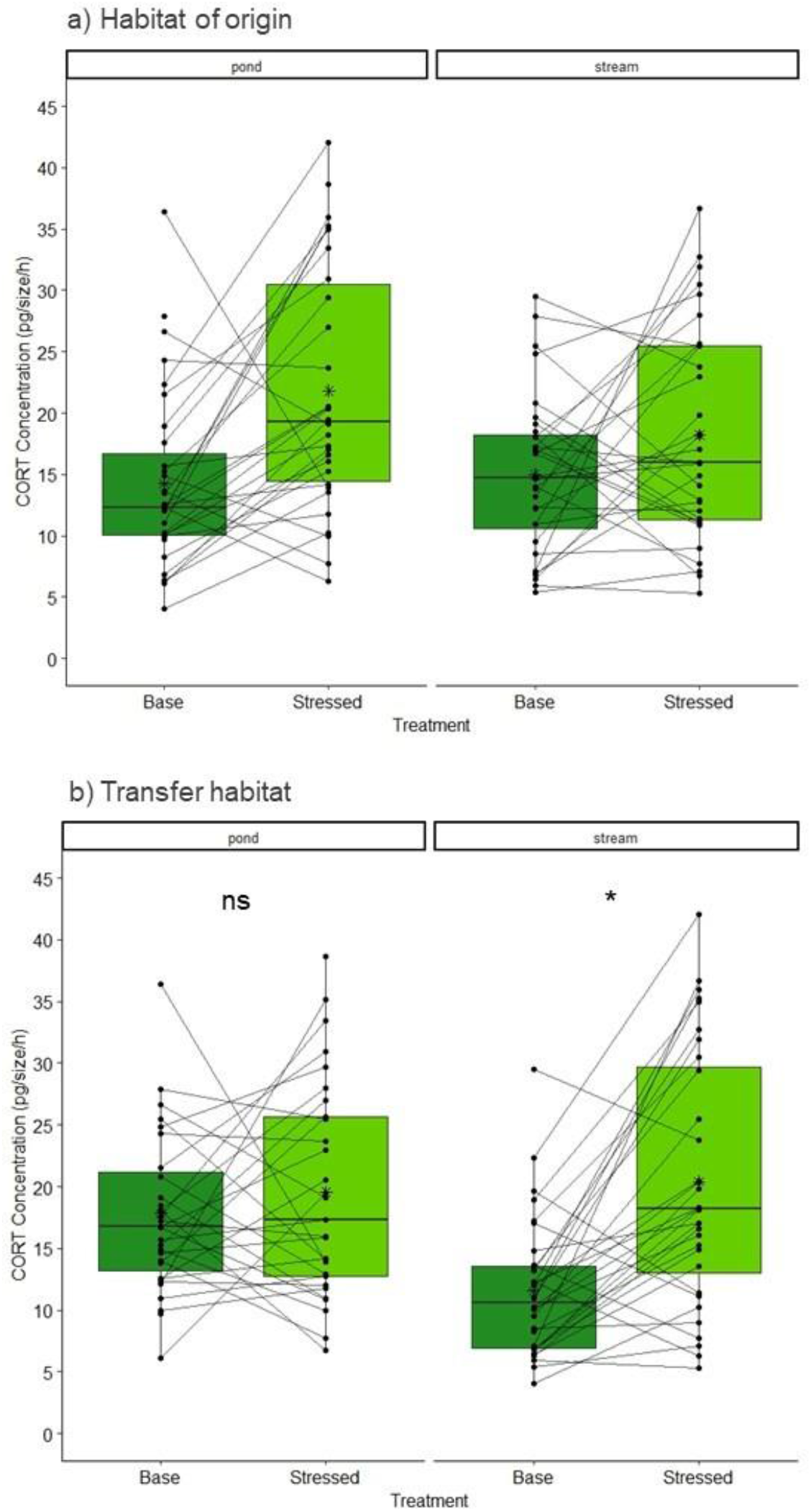
Waterborne corticosterone (CORT) concentration (pg/size/h) in fire salamander larvae (*Salamandra salamandra*). Shown are the baseline (Base) and stress-induced (Stressed) CORT release rates of pond and stream larvae. a) Larvae after the transfer according to their habitat of origin (pond: baseline N = 32, stress-induced N = 32, stream: baseline N = 32, stress-induced N = 31). b) Larvae after the transfer according to their transfer habitat (pond: baseline N = 32, stress-induced N = 31, stream: baseline N = 32, stress-induced N = 31). Each point represents a single CORT measurement, and each line connects the two measurements per individual. The horizontal line represents the median, and the asterisks in the box represent the mean.

**Table 5:** Linear mixed effect model (LMM) for the corticosterone (CORT) release rate regarding the CORT type (baseline or stress-induced) after the transfer, habitat of origin, transfer habitat, the interaction between CORT type and transfer habitat, CORT type and habitat of origin and ID and location. Significant effects are presented in bold.

| CORT release rate (pg/size/h) | Dependent variable |  | Fixed effect |  |
| --- | --- | --- | --- | --- |
|  | Intercept | Estimate | Std Error | p value |
|  | stress-ind. CORT | 0.474 | 0.231 | 0.045 * |
|  | transfer habitat (stream) | -0.901 | 0.484 | 0.185 |
|  | original habitat (stream) | 0.074 | 0.210 | 0.724 |
|  | **Interaction stress-ind. CORT & transfer-habitat (stream)** | **0.777** | **0.267** | **0.005 \*\*** |
|  | **Interaction stress-ind. CORT & original-habitat (stream)** | **-0.558** | **0.267** | **0.041 \*** |

For the transfer habitat pond, we found no difference between baseline and stress-induced CORT (p = 0.390), while we found a difference for the transfer habitat stream (p < 0.05). When comparing the CORT types between the transfer habitats, we found no difference between baseline CORT (p = 0.219) and stress-induced CORT (p = 0.832).

We found no difference in the deviation between baseline and stress-induced CORT in larvae that remained in their original habitat type (St/St & P/P, p = 0.995, figure 5). However, larvae that changed their habitat type differed significantly from each other (P/St & St/P, p = 0.008). Larvae originating from streams and transferred into ponds (St/P) differed significantly from those that remained in streams (St/St, p = 0.048), while we found no difference between larvae that were transferred into streams (P/St) and larvae that remained in streams (St/St, p = 0.906). We found no difference between larvae that were transferred from stream to ponds (St/P) and larvae that remained in ponds (P/P, p = 0.086).

**Figure 5:**
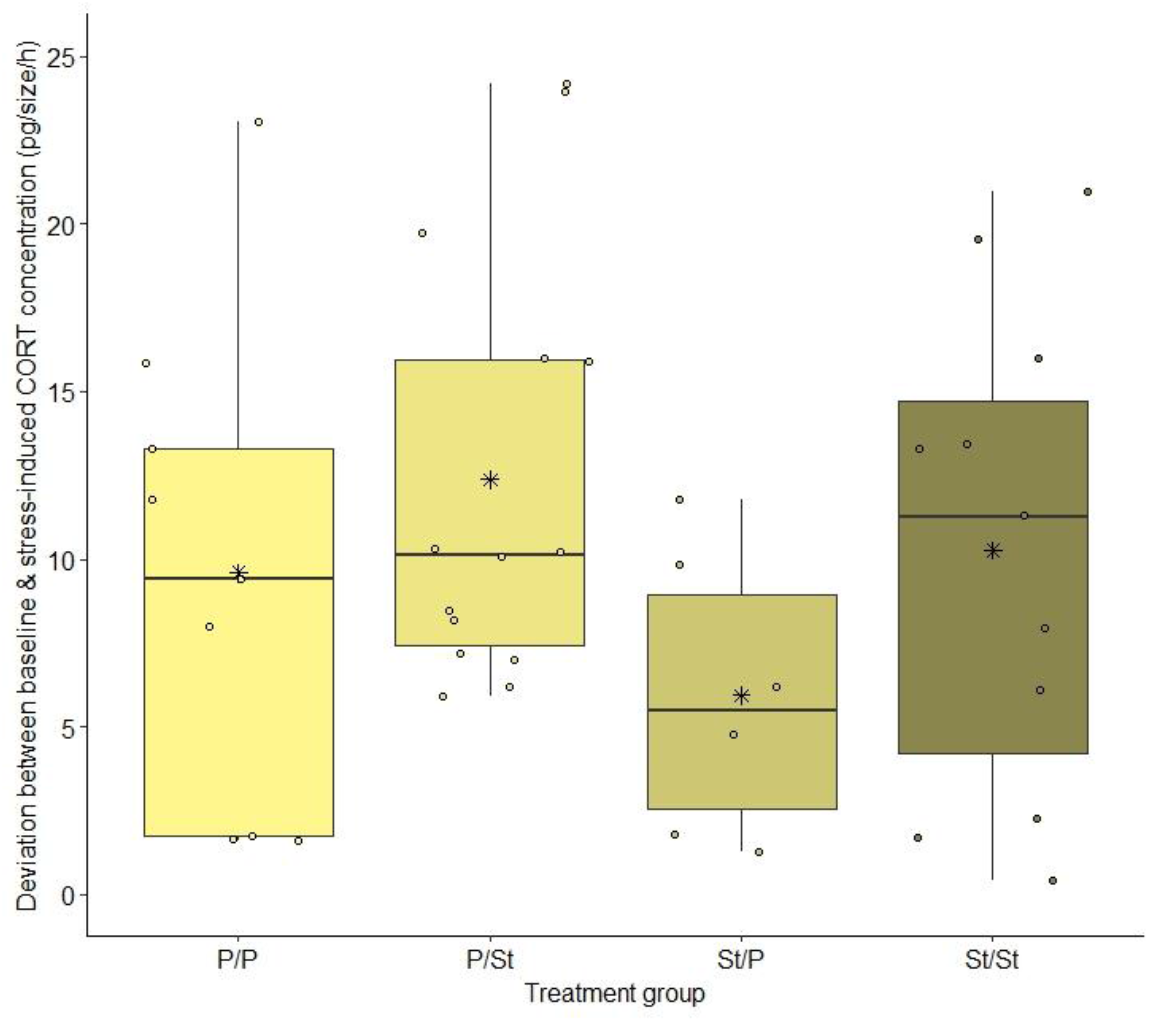
Deviation in baseline and stress-induced CORT concentrations (pg/size/h) between fire salamander larvae from ponds and streams after the transfer for the four treatment groups P/P (pond originated/transferred into ponds, N=15), P/St (pond originated/transferred into streams, N=16), St/P (stream originated/transferred into ponds, N=15) and St/St (stream originated/transferred into streams, N=16). Each point represents the deviation in the CORT concentration of one individual. The horizontal line represents the median, and the asterisks in the box represent the mean.

For the growth rate, we found a significant impact in the transfer group during the transfer period (p = 0.004, Fig. 6). Larvae that originated from streams and remained in streams (St/St) were the only group that significantly differed from all other groups (St/St & P/P: p = 0.019, St/St & P/St: p = 0.006, St/St & St/P: p = 0.037, figure 6). We found no difference in larval growth rate (cm) between larvae transferred into ponds (St/P) or streams (P/St) and larvae that remained in ponds (P/P) (St/P & P/St: p = 0.917, P/St & P/P: p = 0.98, P/P & St/P: p = 0.994).

**Figure 6:**
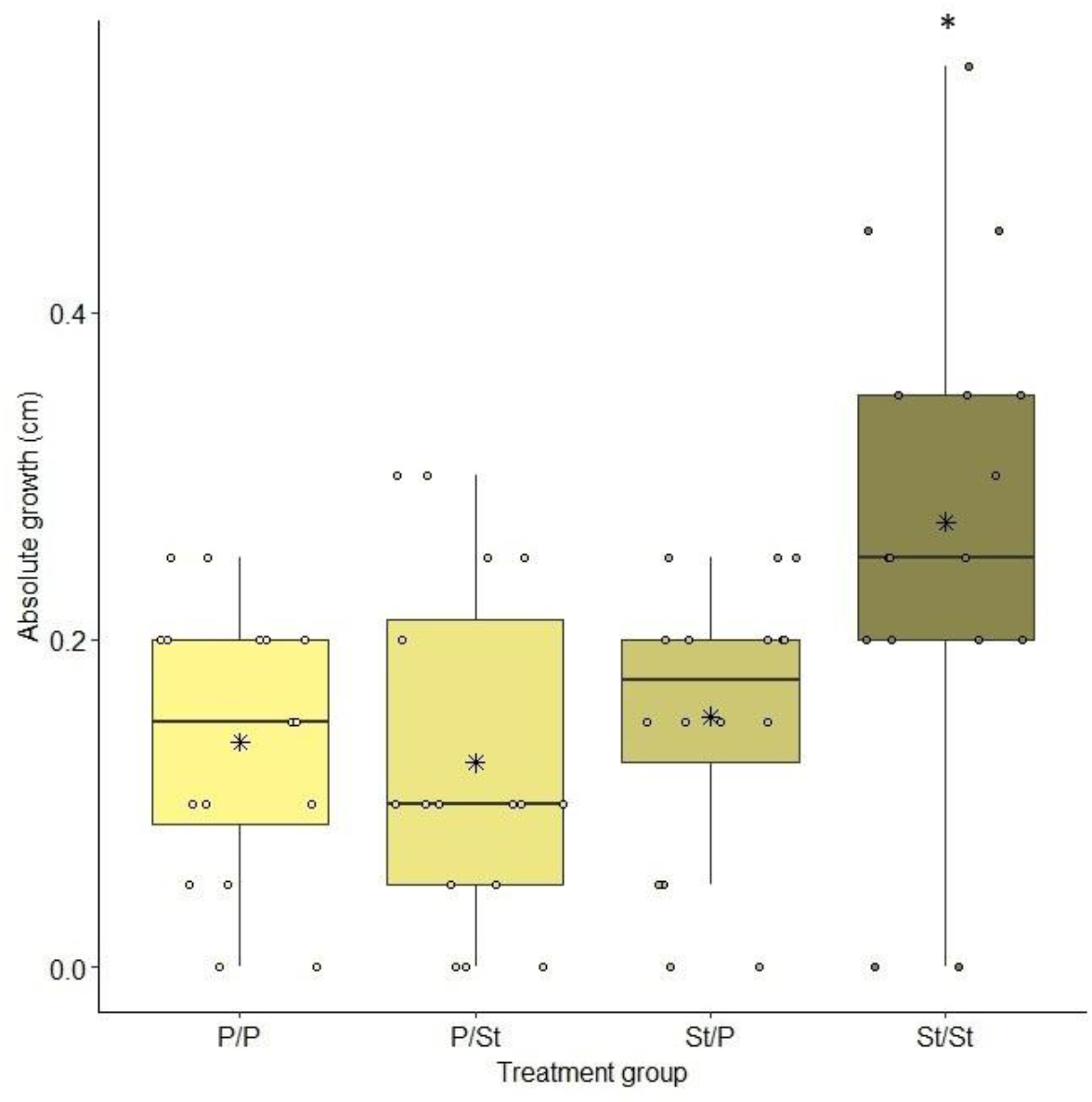
Absolute larval growth (cm) during the transfer period for each treatment group (P/P, P/St, St/P, St/St, each group N=16). Each point represents a single measurement, while the horizontal line represents the median and the asterisks in the box represent the mean.

## Discussion

Our study investigated corticosterone release rates in fire salamander larvae in the field using water-borne hormone sampling. We validated CORT measurements for fire salamander larvae after Gabor and colleagues (2013). Moreover, we focused on the stress that the larvae experience in the different habitat types and whether they can conform to changing habitat conditions.

We showed that pond larvae have significantly higher baseline and stress-induced CORT release rates than stream larvae, confirming our first hypothesis. Ponds are not the typical habitat that fire salamander females choose for larval deposition (Thiesmeier, 2004). In ponds, the water is usually warmer and has less oxygen and food, higher predator abundance and a higher rate of conspecifics than in streams (Bletz et al., 2016; Reinhardt et al., 2013; Weitere et al., 2004). In addition, fire salamander larvae can be cannibalistic, especially when they encounter high densities of conspecifics (Degani, 1993), which is likely to be another stressor for (smaller) larvae. All these factors make ponds the more stressful habitat for fire salamander larvae, which is reflected in their higher CORT release rate.

After the salamander transfer, we found an effect of the original habitat and an even stronger effect of the transfer habitat. Larvae transferred into ponds did not show increases in CORT release compared to the baseline value, whereas larvae transferred into streams showed this increase in stress-induced CORT release. These results indicated that the larvae transferred into streams can react to acute stress, as they showed an increase in corticosterone concentrations after exposure to an acute stressor. We can speculate that this indicates a high potential to conform to the habitat stream. In contrast, the larvae transferred into ponds could not conform to the new habitat, as their stress response indicated signs of chronic stress after two weeks. Chronic stress leads to a downregulation of the stress-induced CORT response towards acute stressors (Dahl et al., 2012; Rich and Romero, 2005), as seen for the larvae that were transferred into ponds. Chronic stress can have severe fitness consequences (Sapolsky, 2002), and a recent study showed that larvae in streams have a higher apparent survival than larvae in ponds (Oswald et al., 2023). Living in a remarkably stressful habitat might be one explanation for this. However, stress may be perceived differently in individuals. Within the treatment groups, we observed strong individual differences in the CORT measurements between baseline and stress-induced CORT. This might reflect individual differences and therefore distinct individualised niches (Müller et al., 2020; Trappes et al., 2022). From our study site, the larvae differ in their genotype according to habitat, with most larvae from ponds belonging to the pond genotype and most larvae from streams belonging to the stream genotype (Hendrix et al., 2017; Steinfartz et al., 2007). However, we did not analyse the genotype of the larvae and thus cannot rule out or prove that the genotype affects individual CORT release rates.

Chronic stress can lead to a decreased size at metamorphosis and influence morphology (Dahl et al., 2012; Kloas et al., 2009; Kulkarni and Gramapurohit, 2017), which might influence survival after metamorphosis. We found that all larvae in ponds and streams were substantially larger after the transfer than before. However, for all larvae that experienced the stressful habitat, the pond, as their habitat of origin, transfer or both, grew less than the larvae that only experienced streams. Increased CORT can negatively affect growth, as found in *Rana temporaria* tadpoles (Ruthsatz et al., 2023) and *Hylarana indica* tadpoles (Kulkarni and Gramapurohit, 2017). In contrast to our findings, a previous reciprocal transplant experiment (Oswald and Caspers, 2022) found a higher growth rate in pond larvae. However, they kept the larvae in the transfer treatment for a longer time (mean of 47 days vs. 14 days in this study). This suggests that growth might be initially faster in stream larvae but is outcompeted with time in pond larvae. Changing abiotic factors over time might also play a role, as Oswald and Caspers (2022) started their transfer experiment in March, while we conducted our study at the end of April. Considering larval physiology is especially important for conservation purposes, the larval stage can be considered the most critical life stage during fire salamander ontogeny. Larval habitats are already heavily affected by the changing climate. Thus, as larvae from both habitat types can conform when transferred into streams, streams should be considered for conservation actions in the future to enhance larval physiological health. Future research should focus on potential physiological differences between pond and stream larvae and the impact that these habitats might have on larval fitness. In addition, this newly validated method can be used to test the influence of other factors on larval physiology, such as pollution, temperature and food availability.

### Conclusion

Fire salamander larvae from ponds and streams differ in several aspects, such as their morphology and behaviour (Oswald et al., 2020; Sabino-Pinto et al., 2019; Schulte, 2008). However, less is known about potential differences in physiological health between larvae from different habitats. We demonstrated that larvae in ponds have a higher baseline and stress-induced CORT release rate than larvae in streams. However, larvae transferred into ponds showed a downregulated stress response towards an acute stressor, while larvae transferred into streams did not. These results indicated (i) that ponds are a more stressful environment for fire salamander larvae and that (ii) larvae can partially conform to a given habitat. In addition, with this study, we validated the use of water-borne corticosterone for fire salamander larvae and thus enabled future use from different perspectives that might influence larval physiology.

## Data availability

Data and code can be found here doi: 10.4119/unibi/2981647.

## Author Contribution

BAC, SK & LS designed the study, LS, PO & MM took the hormone samples in the field, EO, SaK & SK validated the hormone assay, EO and SaK performed the hormone analysis, LS &

PO analysed the data, and LS created the first draft of the manuscript. All authors contributed to the final manuscript.

## Acknowledgement

We thank the Behavioural Ecology group from Bielefeld University for feedback on the statistical analysis and illustration.

