## Supplements for "Stress response of fire salamander larvae differs between habitat types"

**Table 1: Linearity/Parallelism: To assess linearity, we ran a serial dilution of three pooled samples in duplicate.**

| | expected (pg/ml) | measured (pg/ml) | linearity (%) |
|---|---|---|---|
| **sample 1** | | | |
| neat | | 339 | |
| 1:2 | 169.5 | 138 | 81.4 |
| 1:4 | 84.8 | 76 | 89.6 |
| 1:8 | 42.4 | 27 | 63.7 |
| **sample 2** | | | |
| neat | | 393 | |
| 1:2 | 196.5 | 183 | 93.1 |
| 1:4 | 98.3 | 104 | 105.8 |
| 1:8 | 49.1 | 47 | 95.7 |
| 1:16 | 24.6 | 16 | 65.0 |
| **sample 3** | | | |
| neat | | 569 | |
| 1:2 | 284.5 | 280 | 98.4 |
| 1:4 | 142.3 | 130 | 91.4 |
| 1:8 | 71.1 | 57 | 80.1 |
| 1:16 | 35.5 | 23 | 64.8 |

**Table 2: To assess the recovery rate, three pooled samples of different concentrations were spiked with different amounts of corticosterone and measured in duplicate.**

| | expected (pg/ml) | measured (pg/ml) | recovery rate (%) |
|---|---|---|---|
| **sample 1** | | | |
| nondiluted | | 243 | |
| diluted 1:2 | 122 | 112 | 91.8% |
| + 128 pg/ml | 186 (122+64) | 185 | 99.0% |
| + 320 pg/ml | 282 (122+160) | 296 | 103.9 |
| + 800 pg/ml | 522 (122+400) | 598 | 114.6 |
| **sample 2** | | | |
| nondiluted | | 404 | |

| | | | |
|---|---|---|---|
| diluted 1:2 | 202 | 183 | 90.5% |
| + 128 pg/ml | 266 (202+64) | 241 | 90.6% |
| + 320 pg/ml | 362 (202+160) | 412 | 113.8% |
| + 800 pg/ml | 602 (202+400) | 658 | 109.3% |
| **sample 3** | | | |
| nondiluted | | 569 | |
| diluted 1:2 | 285 | 244 | 85.6% |
| + 128 pg/ml | 349 (285+64) | 331 | 94.8% |
| + 320 pg/ml | 445 (285+160) | 481 | 108.1% |
| + 800 pg/ml | 685 (285+400) | 762 | 111.2% |

**Table 3: To assess the recovery rate, corticosterone solutions were added to tap water and measured in duplicate.**

| amounts added to 40 ml tap water (pg) | expected (pg/ml) | measured (pg/ml) | recovery rate (%) |
|---|---|---|---|
| standard 1: 200 | 800 | 839 | 105 |
| standard 2: 100 | 400 | 477 | 119 |
| standard 3: 50 | 200 | 186 | 93 |
| standard 4: 25 | 100 | 109 | 109 |
| std 1 + pool (200+59) | 1036 | 1288 | 124 |
| std 2 + pool (100+59) | 636 | 621 | 98 |
| std 3 + pool (50+59) | 436 | 444 | 102 |
| std 4 + pool (25+59) | 336 | 402 | 120 |

Expected: Samples were resuspended in 250 μl ELISA buffer; thus, the expected values must be extrapolated to ml.
